# Functional Performance of Electrospun Tissue-Engineered Heart Valve Scaffolds Across Geometries Under Aortic Conditions

**DOI:** 10.64898/2026.10.13.758926

**Authors:** Hannes Dillinger, Aref Saberi, Mattia Manenti, Sil Gussenhoven, Iva van der Burgh-Vrees, Carlijn V.C. Bouten

## Abstract

Electrospun, bioresorbable polymer scaffolds are a promising platform for tissue-engineered heart valves (TEHVs) owing to their fibrous microstructure, tunable mechanics and capacity to support host-cell infiltration and remodeling. Recent in-vitro parametric studies of bioprosthetic heart valve (BHV) geometry have used silicone cast valves to link design parameters to hemodynamic performance and leaflet strain, but the functional behavior of electrospun whole-valve TEHV scaffolds across distinct leaflet geometries and clinically relevant aortic conditions remains poorly characterized. Here, we present an end-to-end workflow in which polycaprolactone (PCL) is electrospun directly onto parametrized, 3D-printed mandrels derived from two established analytical leaflet geometry models, yielding 21 whole-valve scaffolds across four geometries without cutting and assembling flat fiber sheets or suturing of flat fiber sheets. Fiber diameter, orientation and pore area were assessed by scanning electron microscopy and found to be consistent across scaffolds and geometries. Two scaffolds of each of four geometries were tested in a pulse duplicator under hypotensive, normotensive and hypertensive aortic conditions. All scaffolds opened and closed physiologically under all conditions, with a mean maximum geometric orifice area (GOA) of 44.5% of the valve area and a mean peak transvalvular pressure gradient of 10mmHg. Leaflet geometry had no apparent systematic effect on flow and pressure waveforms or on GOA. The mean regurgitant fraction was 82%, most likely due to leakage through the porous fiber network rather than incomplete leaflet closure. Electrospun whole-valve scaffolds thus achieve favorable leaflet kinematics at clinically relevant aortic conditions and the presented workflow provides a reproducible, parametric framework for their fabrication and translation toward regenerative heart valve replacement.

## Introduction

Heart valve disease is a major global health burden, and despite decades of engineering progress, no prosthetic design fully replicates the biological, mechanical and regenerative capacity of a native valve. Tissue-engineered heart valves (TEHVs) address this gap as living valve substitutes capable of growth, self-repair and physiological remodeling in response to hemodynamic stimuli, properties absent from both mechanical prostheses and conventional bioprosthetic heart valves (BHVs).

The field has shifted toward in-situ tissue engineering, in which cell-free scaffolds are implanted directly and subsequently repopulated by host cells, avoiding the complexity, cost and regulatory burden of bioreactor-based cell seeding Stassen et al. (2017); Motta et al. (2018); Fioretta et al. (2021). Transcatheter valve replacement techniques have shaped TEHV design requirements from an early stage Motta et al. (2018); Cheung et al. (2015); Xiling et al. (2022). Current TEHV concepts Fioretta et al. (2021); Cabrera et al. (2017) are designed as stented constructs integrated into expandable frames for transcatheter delivery in line with the corresponding valve geometry. Bioresorbable stent materials are increasingly used, since they provide temporary mechanical support without permanently scaffolding the vessel wall, consistent with the regenerative goals of TEHVs.

In parallel, advances in transcatheter aortic valve replacement (TAVR) technologies have highlighted the importance of scalable manufacturing, simplified deployment and cost reduction to broaden access to heart valve replacement. Appa et al. (2022) demonstrated a balloon-expandable TAVR platform incorporating either bioprosthetic or polymeric leaflets, with a design intended to facilitate deployment without rapid pacing and to enable anchorage even in the absence of calcification. The development of multiple valve sizes, compatibility with clinically established crimping dimensions and favorable hemodynamic and fatigue performance illustrate how device engineering can support broader clinical applicability. Such approaches are particularly relevant to expanding access in low- and middle-income countries, where the burden of rheumatic heart disease remains substantial and the cost and availability of conventional valve replacement limit treatment options. However, while these advances demonstrate progress toward more accessible transcatheter valve technologies, scalable production must also address reproducibility, material consistency, manufacturing throughput and quality control. For TEHVs, fabrication strategies that enable reproducible production of complete valve scaffolds while minimizing labor-intensive assembly steps may therefore offer an important route toward cost-effective scale-up and wider clinical translation Motta et al. (2018).

The importance of leaflet geometry for valve function and durability has been studied in conventional BHVs. Recently, Bueno et al. (2026) systematically varied geometric parameters, including leaflet thickness, diameter, belly curvature, free-edge angle and leaflet height, in silicone cast valves modeled on the Trifecta BHV (Abbott) and assessed their influence on hemodynamic performance and leaflet strain using Doppler echocardiography and image correlation. The work established quantitative relationships between BHV design parameters and functional outcomes and showed that in-vitro parametric testing can guide valve design. Silicone cast valves, however, differ from electrospun TEHV scaffolds in material properties, microstructure, porosity and regenerative capabilities. The geometry-function relationships identified for homogeneous silicone leaflets cannot be assumed to hold for fibrous and porous PCL(-BU) scaffolds, whose mechanical behavior depends on the electrospinning process and mandrel geometry. The strain-based failure and durability metrics relevant to BHVs are also not the main design criteria for TEHVs, where in-vivo remodeling and biocompatibility are the primary endpoints.

Manufacturing workflows for electrospun TEHVs Chen et al. (2009); Kluin et al. (2017); Alavi et al. (2017) still rely on cutting flat fiber sheets and assembling individual leaflets, multi-mandrel spinning Capulli et al. (2017) or, as in Motta et al. (2023), on post-spinning embossing to achieve the final valve shape. These multi-step procedures may introduce operator dependence and variability in leaflet shape. Changes in fiber thickness distribution and local mechanical behavior obscure the relationship between scaffold geometry and functional performance, which is critical for reproducible design and clinical translation. Few studies have examined how collector geometry influences the hemodynamic behavior of electrospun whole-valve scaffolds and fewer still have benchmarked such scaffolds across the full range of clinically relevant aortic pressure conditions. Direct spinning on mandrels have resulted in ruptured scaffolds van Lieshout et al. (2006).

Three gaps follow from this literature, each with direct consequences for clinical translation. First, systematic comparisons of mandrel geometries for electrospun whole-valve TEHV fabrication remain limited. Differences in fabrication approaches and the lack of standardized workflows make it difficult to assess how mandrel geometry affects leaflet shape and fiber characteristics across scaffolds. Improved reproducibility could support more consistent quality control and facilitate the development of scalable and cost-effective manufacturing processes. This could enable production available beyond specialized centers, including in low- and middle-income settings. Second, geometry-function relationships in electrospun TEHVs remain poorly characterized through the stage of in-vitro hemodynamic testing under demanding aortic conditions. A cell-free scaffold must provide full valve function from the moment of implantation, before host cells have repopulated and remodeled it. Its initial geometry therefore determines leaflet coaptation, transvalvular gradient and regurgitation in the patient and it also sets the mechanical environment that guides subsequent tissue formation. Without these relationships, scaffold design cannot be rationally adapted to different valve sizes or patient requirements. Third, functional benchmarking of electrospun TEHV scaffolds at clinically relevant aortic pressure conditions is insufficient. Patients requiring valve replacements present with a wide range of hemodynamic states, from hypotensive low-output conditions to the hypertension that is common in this population and a regenerative valve must open fully and close competently across this range. Testing at a single nominal pressure leaves this functional range undefined. To address these gaps, we present a reproducible end-to-end workflow for electrospinning PCL whole-valve scaffolds directly onto parametrized, 3D-printed mandrels and evaluate scaffold microstructure and hemodynamic valve function under hypotensive, normotensive and hypertensive aortic conditions. In contrast to parametric BHV studies using surrogate materials, our approach evaluates the electrospun scaffold itself, preserving the fiber microstructure and mechanical isotropy relevant to in-vivo TEHV performance and remodeling.

## Methods

### Computer-aided valve and holder design

Aortic leaflet geometry varies even among healthy individuals, so reproducible mandrel design requires a simplified mathematical description. We implemented two established models: Hamid et al. (1986) defines a tilted elliptic half-paraboloid, with parameters set iteratively to match a prescribed leaflet surface area (implementation was based on the work of Loerakker et al. (2013)) and Utku Gulbulak et al. (2020) defines leaflet surfaces from commissure and belly splines specified by point coordinates (see Figure 1). For the latter, a variant corresponding to the coordinate transformation of the commissure curve based on the axis coordinate was implemented. The generated meshes are illustrated in Figure S1. We selected these geometries to allow direct comparison with extensive computational studies of TEHV remodeling and diastolic stress distributions by Loerakker et al. (2013); Visser et al. (2025). Mandrel radius and height were set to 13.5 mm and 18 mm, corresponding to a healthy adult aortic valve inner diameter of 27 mm. Valve holders for functionality testing were generated automatically from each mandrel geometry to fix the valve base while allowing free leaflet motion, so that hemodynamic results reflect scaffold geometry rather than interaction with the holder.

**Figure 1.**
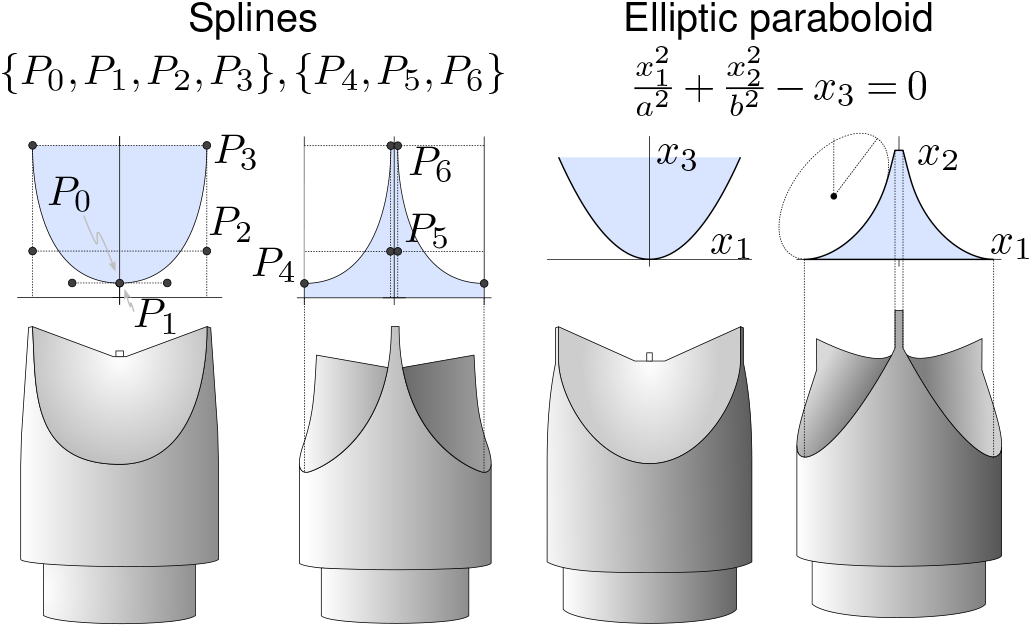
The model by Utku Gulbulak et al. (2020) (left) is defined by points *P*_0_ to *P*_3_ across and inside the leaflet, respectively, which are used to define smooth splines. The model by Hamid et al. (1986) (right) is defined by a tilted elliptic paraboloid. Here, the projection onto the cylindric geometric was performed by setting the coordinate *x*_1_ equal to the arc’s length.

The 3D models were exported in stereolithography (STL) format and 3D printed using polyactic acid (X1 Carbon, Bambu Lab USA Inc, Austin TX, USA), fixed on existing target cylinders and coated with conductive spray paint (843AR SUPER SHIELD, MG Chemicals, Ontario, Canada). The target cylinders were covered with aluminum foil below the 3D printed geometry.

### Electrospinning and scaffold preparation

PCL (Purasorb PC 12, Corbion) was dissolved in 9:1 (v/v) chloroform:methanol at 17.45% (w/w) and stirred overnight. Electrospinning was performed on a commercial machine (CLI-751, Vivolta) at 30% relative humidity and 23°C, with needle and mandrel potentials of 22 kV and −1 kV, respectively. Precursor was supplied at 20 µL min^−1^ through a 0.8 mm diameter needle; the target was placed 20 cm from the needle and rotated at 300 rpm. Spinning began at the aluminum foil base below the mandrel, advanced over the mandrel across a 35 mm stroke at 15 mm s^−1^ and stopped once thickness at the valve base, monitored continuously by a laser measurement, reached 450 µm (about 1 hour per scaffold). Scaffolds dried overnight on the mandrel, were dipped in 70% ethanol to allow for fiber swelling and released with pressurized air (see Figure 2). The scaffolds were trimmed at the commissure tips, effectively removing the excess height caused by the spinning process. Scaffolds were sutured to holders at the commissures with 5-0 monofilament (Ethilon, Ethicon, Cincinnati OH, USA) and fixed at the base with an o-ring.

**Figure 2.**
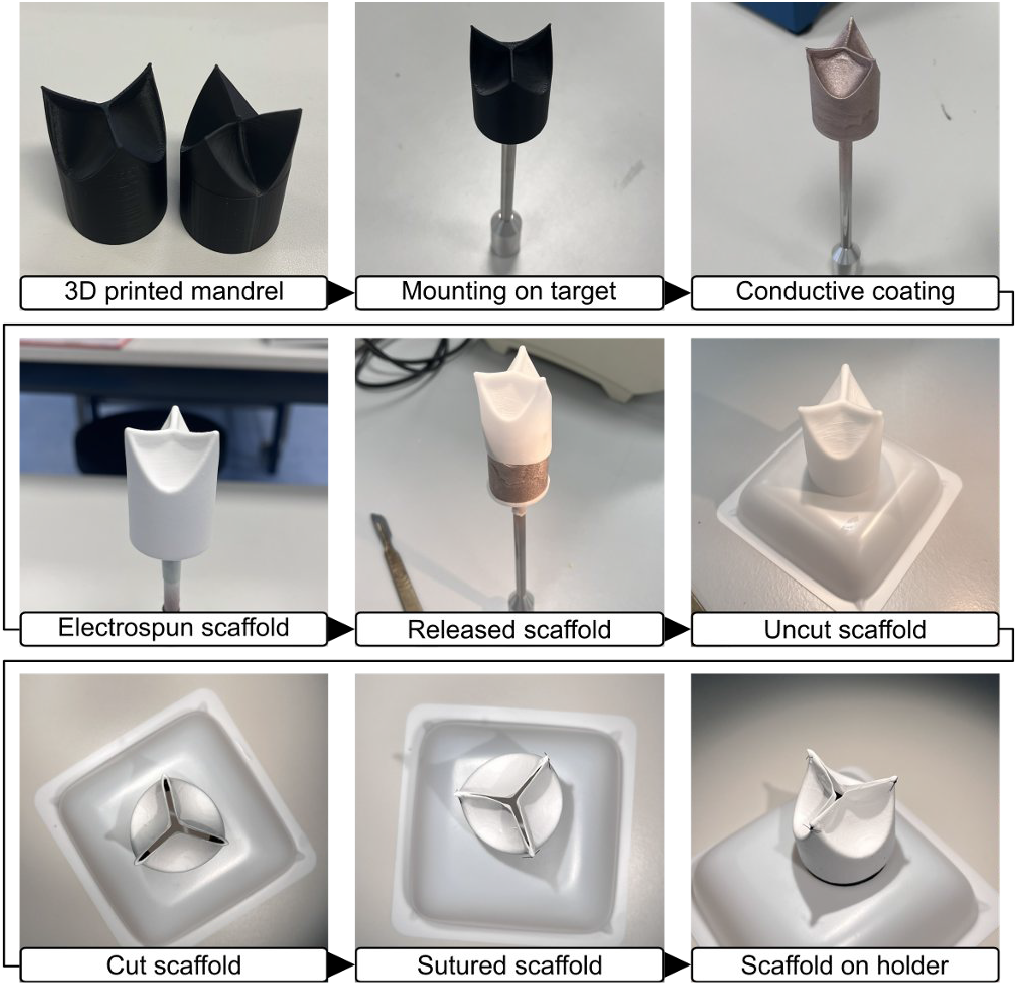
Complete workflow for scaffold manufacturing. 3D printed mandrels were mounted on existing targets, conductively coated and used for electrospinning. After drying, the scaffold was released using pressurized air and cut at the top. The scaffold was sutured to its holder using surgeon’s knots at the commissures.

### Microstructural characterization

Aluminum foil samples collected during spinning were sputter-coated with gold (JFC-2300HR, Jeol, Tokyo, Japan) to prevent charging and imaged by scanning electron microscopy (SEM; JCM-7000 NeoScope, Jeol). Fiber diameter, orientation and pore area, which influence cell attachment, migration and ingrowth, were quantified using DiameterJ (Hotaling et al., 2015) and visualized with quanfima (Shkarin et al., 2019). To assess whether foil samples are representative of the leaflet microstructure, one scaffold was sacrificed after functionality testing and SEM samples were taken from the leaflet center, leaflet top and the aluminum foil.

### Functionality testing

Hemodynamic performance was assessed in a commercial pulse duplicator (HDT-500, BDC Laboratories, Wheat Ridge CO, USA) configured for aortic testing at hypotensive (100*/*60 mmHg), normotensive (120*/*80 mmHg) and hypertensive (160*/*100 mmHg) conditions, covering the clinically relevant pressure range from borderline low to pathologically elevated aortic pressures. Stroke volume was set to 60 mL and 80 mL for hypo-/normo- and hypertensive testing, respectively. No ventricular compliance was used. The pump was operated at 60 min^−1^ with waveform profile of 33 % of systole, yielding a physiologically skewed sinusoidal waveform with an early systolic peak. Purified water at 21 °C was used as the working fluid. Flow was measured by an ultrasonic sensor (ME25PXN319, Transonic, Ithaca NY, USA) positioned upstream of the valve and connected to a flowmeter (TS410, Transonic, Ithaca NY, USA) which was used to export flow curves and calculate the regurgitant fraction. Valve opening and closing behavior and the geometric orifice area (GOA) were quantified from high-speed video recordings (a2A2590-60ucPRO, Basler, Ahrensburg, Germany, 800×800 pixels, up to 150 fps) through the aortic compliance chamber inspection window, using a custom Python segmentation script. GOA was expressed as a percentage of the valve area to allow comparison across geometries and sizes. This direct optical GOA measurement complements the ISO 5840-defined effective orifice area (EOA) which was not evaluated here. Because camera and pulse duplicator were not synchronized, flow and GOA data were aligned by matching the start of forward flow (FFstart) to the rising edge of the GOA (see Limitations section).

## Results

### Successful fabrication of homogeneous scaffolds and porous fiber morphology

Four mandrel geometries were generated from the Hamid and Gulbulak models and all mandrels were successfully 3D-printed with a coaptation zone thickness of 1 mm. A total of 21 whole-valve scaffolds were electrospun and released from the mandrels without damage to the fiber network (Figure 2). Eight scaffolds, two of each of the four geometries selected for functional testing, are shown in Figure 3 and were evaluated in the remainder of the study. Comparison of SEM samples from the leaflet body and the aluminum foil of the sacrificed scaffold reveals comparable fiber diameter and orientation distributions as shown in Figure 4. This supports aluminum foil-based SEM as a practical quality control strategy that does not require sacrificing functional scaffold material and is an important feature for a scalable workflow. Subsequent SEM results were derived from aluminum samples.

**Figure 3.**
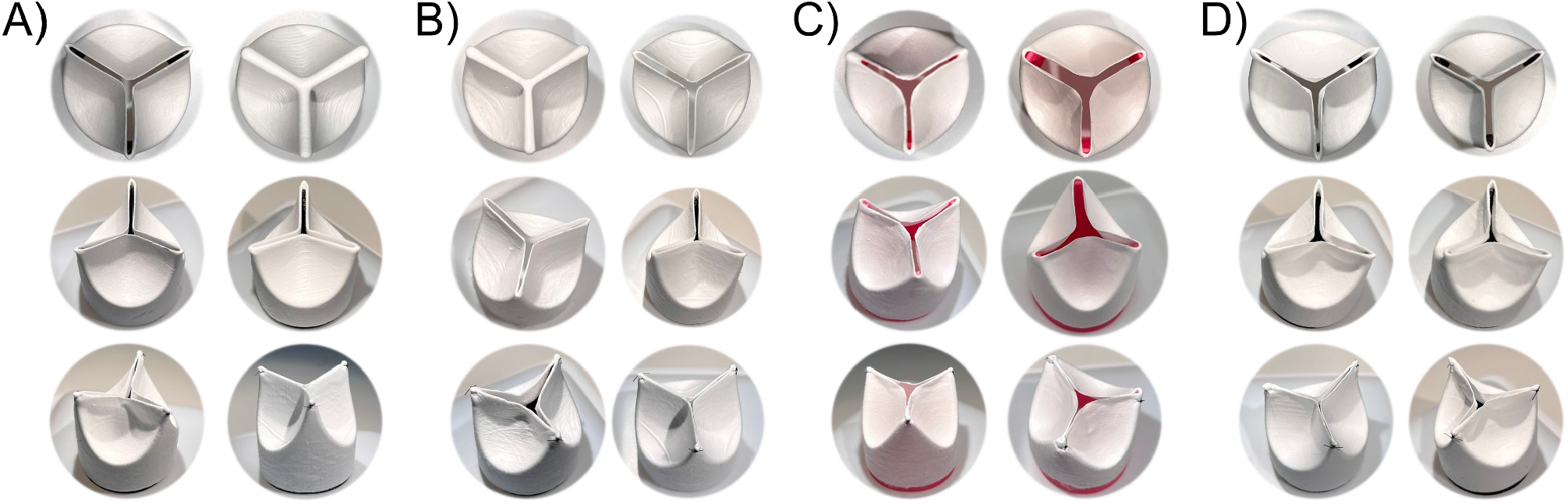
Photographs of the manufactured scaffolds of geometries A) Gulbulak *D* = 0, B) Gulbulak *D* = 0.5, C) Hamid *A* = 2.8, *B* = 1.4, D) Hamid *A* = 4.0, *B* = 2.4. First and second rows show top and side view of the cut valve, third row shows sutured valve. For two scaffolds from A) and B), no cut top view photograph was available.

**Figure 4.**
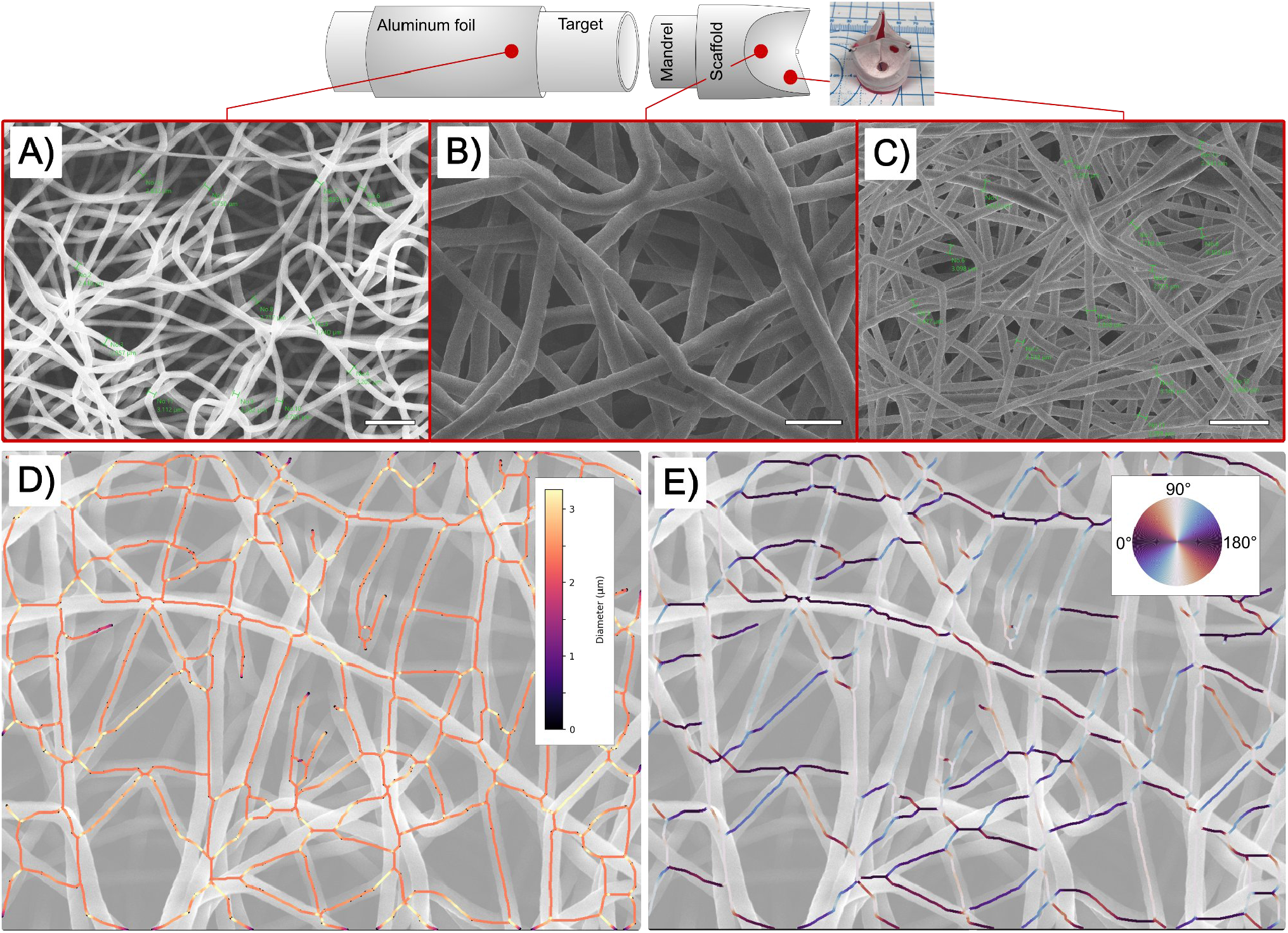
SEM images showing qualitative agreement between A) the aluminum foil sample and B), C) leaflet samples (center and top) at different magnifications (scale bars 20 µm, 10 µm, 20 µm respectively). Automated fiber analysis of scaffold #1.7: D) fiber diameters are homogeneous, of mean diameter 2.5 µm and E) fiber orientations show no preferred direction. Vertical fibers are rendered in light colors, which may limit their visibility.

SEM evaluation in Figure 4 and Figure S2 confirms qualitatively homogeneous fiber morphology across the geometry types, with smooth individual fiber surfaces and no merging of crossing fibers. Quantitative analysis across all scaffolds (incl. the subset for functional testing) yields mean fiber diameters of 3.3 ± 1.3 µm and mean pore areas of 7.3 ± 5.2 µm^2^, consistent with dimensions reported to support cellular infiltration and ECM deposition in electrospun

TEHV scaffolds Wissing et al. (2019). Fiber orientation is randomly distributed, indicating the absence of systematic alignment artifacts from the spinning process. Detailed fiber analysis can be found in Supplementary Figure S3 and Figure S2.

### Functional performance across geometries and aortic conditions

All functionally tested electrospun whole-valve scaffolds demonstrated physiological opening and closing behavior across the full range of aortic pressure conditions tested, without flow obstruction or leaflet prolapse. Video snapshots are shown in Figure 5 and slow-motion video recordings can be found in the Supplementary Material section.

**Figure 5.**
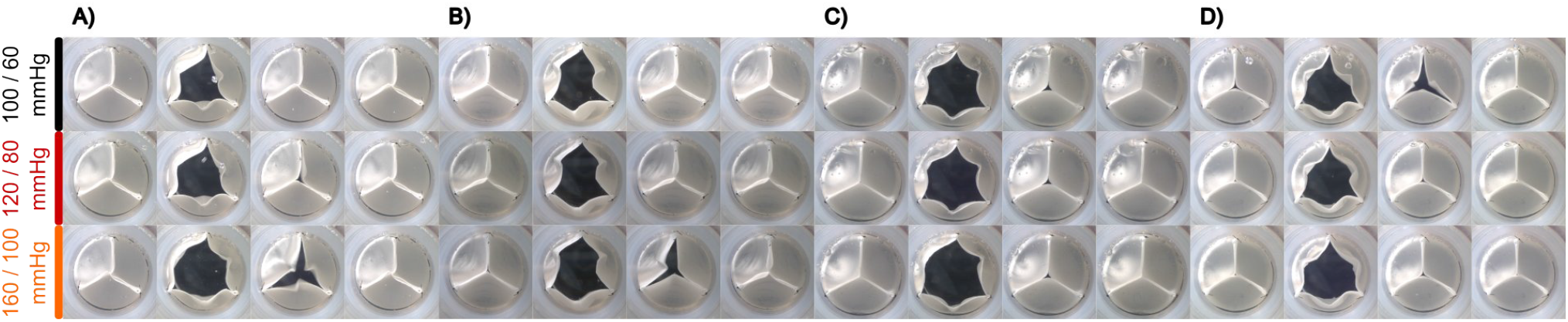
Camera snapshots at *t* = {0.1, 0.3, 0.5, 0.7} for A) Gulbulak *D* = 0, B) Gulbulak *D* = 0.5, C) Hamid *A* = 2.8, *B* = 1.4, D) Hamid *A* = 4.0, *B* = 2.4 at hypo- (100*/*60 mmHg), normo- (120*/*80 mmHg) and hypertensive (160*/*100 mmHg) conditions.

Cycle-to-cycle variability over 10 consecutive cycles was negligible for flow, pressure and GOA. All line plots in the current manuscript include a *±*2*σ* band which is barely visible due to the small variance.

Across the four mandrel geometries, flow and pressure waveforms shown in Figure 6 and Supplementary Figure S4 were similar, with differences concentrated in the backflow phase. Flow maximum was 30 L min^−1^ to 34 L min^−1^ at hypertensive condition. The closing volume across geometries and pressure conditions was comparable (hypotensive 1.50 ± 0.26 mL, normotensive 1.44 ± 0.15 mL, hypertensive 1.75 ± 0.24 mL). The leakage volume increased with increasing pressure, reaching a maximum of 64.80 mL for a stroke volume of 89.82 mL at hypertensive conditions (see Figure 6, C.1).

**Figure 6.**
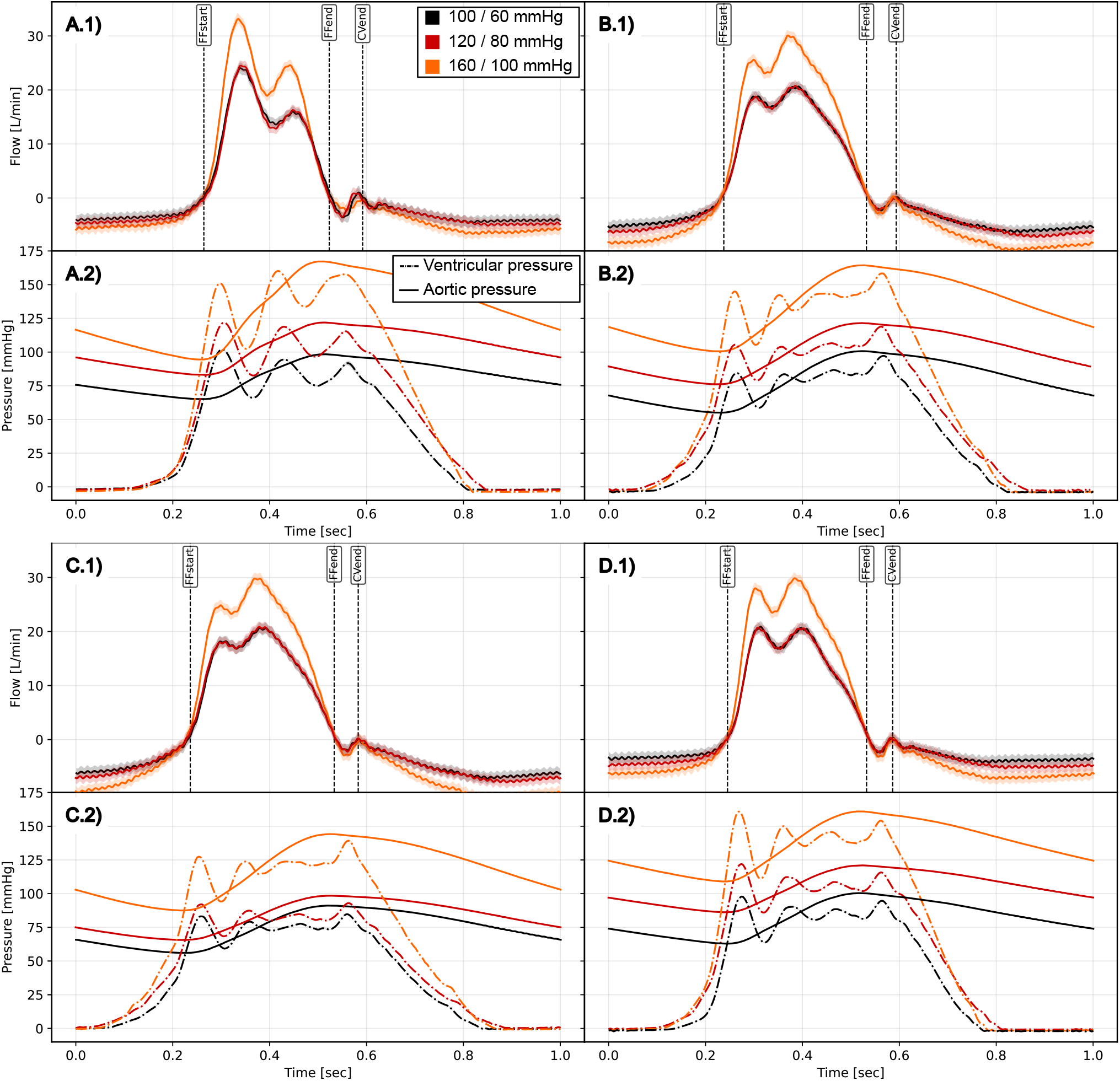
Flow .1) and pressure curves .2) for geometries A) Gulbulak *D* = 0, B) Gulbulak *D* = 0.5, C) Hamid *A* = 2.8, *B* = 1.4, D) Hamid *A* = 4.0, *B* = 2.4. Line plots include *±*2*σ* of the same color. The marker positions for forward flow start (FFstart), forward flow end (FFend) and closing volume end (CVend) are given for the hypotensive case (black line).

For the Gulbulak *D* = 0 model scaffold, RMS forward flow was 250.51 L s^−1^, 250.73 L s^−1^ and 348.17 L s^−1^ for hypo-, normo- and hypertensive conditions, respectively and closing volumes were consistently low (1.39 *−* 1.78 mL). The mean regurgitant fraction of the scaffolds was 82 %.

The opening and closing characteristics were physiological while exhibiting no signs of wear and tear during 15 min of testing. Camera snapshots in Figure 5 show comparable opening and closing over time, yet the scaffold’s orifice shape during peak systole was different between the geometries and within hypo-, normotensive and hypertensive pressure conditions. The maximum GOA across all scaffolds was 62 %, 73 % and 76 % at maximum opening under hypo-, normo- and hypertensive conditions and returned to zero during diastole, suggesting no visible residual orifice. The mean of the maximum GOA for all scaffolds was 44.51 % and the mean maximum transvalvular pressure gradient was 10 mmHg, indicating favorable hydrodynamic resistance.

For the Hamid *A* = 2.8, *B* = 1.4 model, the temporal evolution of the GOA is shown in Figure 7. The GOA waveform exhibited a rapid opening phase followed by a plateau. The valve was opened and closed for 30% for 70% of the cycle time period, respectively, which was considered physiological. For hypo- and normotensive conditions, the GOA waveform was identical. For hypertensive conditions, the GOA increased by 10 %.

**Figure 7.**
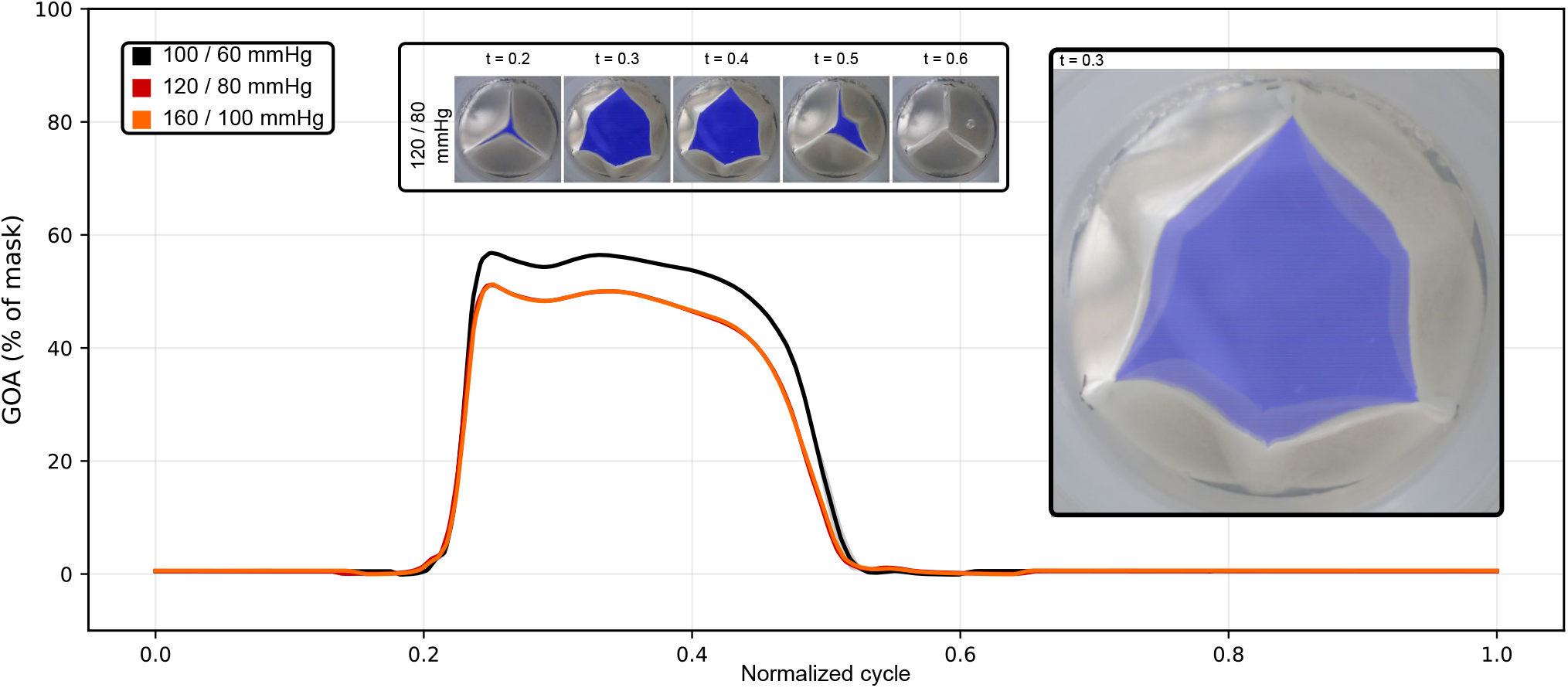
Geometric orifice area (GOA) over normalized cycles at {100*/*60, 120*/*80, 160*/*100} mmHg for the Hamid *A* = 2.8, *B* = 1.4 model. GOA data extracted from video recordings has been averaged over 14 consecutive cycles (14 s) and mean (line) and standard deviation (*±*2 SD, bands) are shown but not visible due to their negligible size. The valve showed physiological movement over the cycle and closed completely. The valve’s movement did not change between subsequent cycles and hypo- and normotensive conditions which used the same stroke volume. For hypertensive conditions, however, the GOA increased by 10 %, most probably because the stroke volume was increased by 33 % to match the set pressure condition. Insets show the scaffold at different time points *t* in the cycle and an overlay of the scaffold at *t* = 0.3 at the different pressure conditions.

To investigate the relationship between GOA and flow, phase plots are shown in Figure 8. The scaffolds exhibited a hysteresis between the opening and closing paths. During the maximum opening phase, oscillation in the flow caused a loop in phase plot (cf. Figure 6). Closing of the scaffolds is comparable between the pressure conditions. All tested scaffolds exhibited similar characteristics in the phase plots (cf. Figure S5).

**Figure 8.**
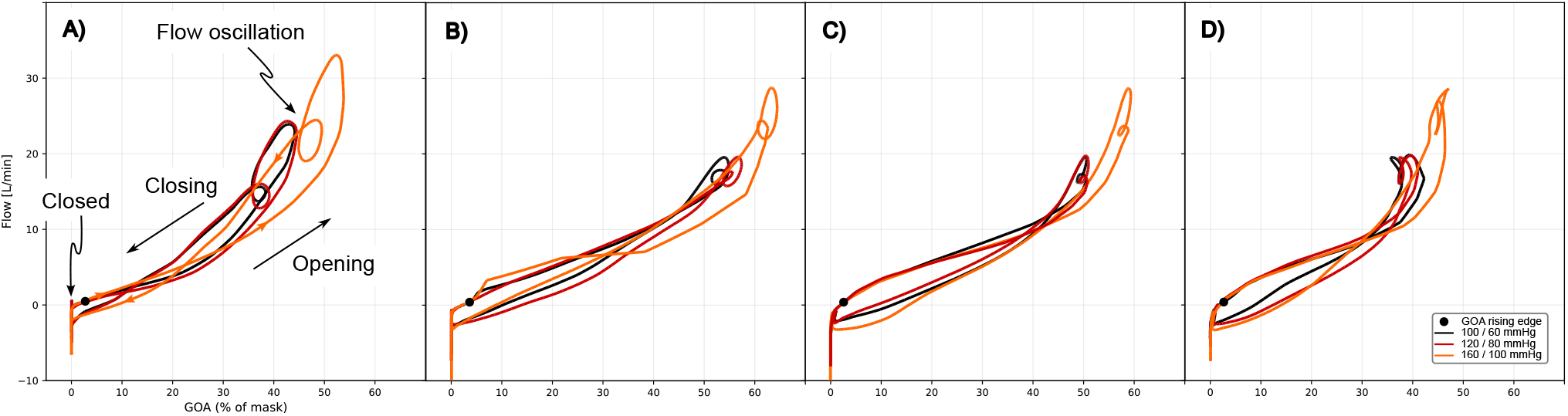
Phase plots GOA vs. flow for the different geometries. A) Gulbulak *D* = 0, B) Gulbulak *D* = 0.5, C) Hamid *A* = 2.8, *B* = 1.4, D) Hamid *A* = 4.0, *B* = 2.4. Hypo- and normotensive curves are comparable while hypertensive curves exhibit increased flow and GOA values. This is to be expected since the stroke volume was increased for the hypertensive case and the opening of the valves is flow-driven. The closing of the valve, however, is also comparable between all three pressure conditions.

## Discussion

Electrospinning is the most widely used method for fabricating TEHV scaffolds, but systematic functional evaluation of electrospun whole-valve constructs under clinically relevant aortic conditions remains limited. Valve geometry is a well-established determinant of hemodynamic performance in conventional BHVs. Bueno et al. (2026) showed that parametric variation of silicone cast BHV geometry, including leaflet thickness, diameter, curvature and free-edge angle, changes hemodynamic performance and leaflet strain under controlled in-vitro conditions. That study offers a useful methodological framework and indicates that geometry is a primary determinant of valve function, but its findings apply to homogeneous, isotropic silicone constructs designed to model BHV durability. The present study addresses a different question: how mandrel geometry influences the hemodynamic function of electrospun, fibrous TEHV scaffolds, where geometry and microstructure are inseparable and set by the fabrication process itself and where the goal is regenerative remodeling capacity rather than structural durability.

A central contribution of this work is that direct electrospinning onto 3D-printed mandrels reproducibly yields whole-valve PCL scaffolds across multiple relevant geometries without cutting, suturing of flat sheets or embossing.

In total, 21 whole-valve scaffolds were electrospun and released from the mandrel without damage to the fiber network. This approach avoids post-spinning manufacturing steps, reduces operator dependence and may preserve the direct link between mandrel geometry and the resulting scaffold microstructure and mechanics. Mold-based geometry variation casts the same material into different shapes; altering the spinning mandrel, in contrast, may change both the leaflet geometry and the deposition of the fiber network. Each geometric variant is therefore an independent experimental condition rather than a different mold for the same material. The parametric design pipeline is automated and openly available, which should make it straightforward to run further parameter studies linking mandrel shape to functional outcomes and to compare these results directly with computational TEHV simulations.

Process reproducibility for electrospun TEHV scaffolds has been addressed previously, in particular during the development of fully polymeric scaffolds for in-situ TEHVs Putti et al. (2015); Simonet et al. (2014). Here, we adapt the workflow to PCL, a widely available, low-cost and extensively characterized precursor material. Previous work showed comparable in-vitro results for PCL and for PCL-BU / PC-BU Dewy C. van der Valk, Charlotte M. Hoes, Yunia M. H. Rasenberg, Paul A. A. Bartels, Livia Angeloni, Bente J. de Kort, Paul J. F. M. Janssen, Frank P. T. Baaijens (2025), the latter being used in-vivo. The present study addresses fiber morphology and acute hemodynamic function rather than degradation, remodeling or long-term durability, so the limitations of PCL for in-vivo use do not affect the comparisons made here. Using PCL lowers material cost and the barrier to entry for parametric studies, which would be costly with implant-grade polymers of limited availability. Process reproducibility therefore serves here as a prerequisite rather than as the contribution itself: it allows mandrel geometry to be varied as the only parameter and its effect on valve hemodynamics to be quantified.

Microstructural characterization of a subset of eight scaffolds showed that fiber morphology was consistent across the four geometries, where the aluminum foil samples were evaluated. Mean fiber diameters fall within ranges associated with favorable cell adhesion and infiltration in electrospun TEHV scaffolds. These properties have no direct analog in silicone BHV surrogates, but they are directly relevant to the regenerative performance of an implanted TEHV. Fiber characteristics measured on aluminum foil samples agreed with those of the sacrificed leaflets, which supports aluminum foil as a non-destructive quality control method. This may be useful for maintaining throughput in a manufacturing pipeline aimed at scalable TEHV production in the future.

Functionally, the evaluated electrospun whole-valve PCL scaffolds showed physiological opening and closing across the full range of hypotensive to hypertensive aortic conditions. This result is not self-evident for porous fibrous scaffolds. Unlike other materials used in BHV parametric studies, electrospun PCL leaflets are permeable to water, they may be mechanically anisotropic at the fiber level and subject to swelling and fiber rearrangement under flow, which narrows the gap between experiment and application significantly. The elevated regurgitation of the uncoated scaffolds investigated here may be direct a consequence of scaffold porosity and the low viscosity of water relative to blood, rather than a failure of leaflet coaptation. Complete GOA closure during diastole supports this interpretation. This important finding supports the translational evaluation: in-vivo blood clotting and fibrin deposition within the scaffold pores are expected to substantially reduce leakage upon implantation Motta et al. (2023). In BHV parametric evaluation using impermeable silicone, this mechanism is absent and regurgitation reflects only leaflet coaptation geometry.

The flow and pressure characteristics over full cycles did not differ substantially between scaffold geometries. During forward flow, this is to be expected since the flow profile is dictated by the pump’s piston motion (“flow-driven”). In late systole and diastole, however, the flow deceleration causes an adverse pressure gradient and the morphology and material properties of the root and valve leaflets define flow vortices and pressures (“pressure-driven”). In contrast to other works, we reported detailed flow and pressure waveforms over time. Together with the pump settings, flow and pressure define the boundary conditions for the scaffold’s characteristics in the in-vitro comparisons. For our samples, the pressure waveforms did not change with the scaffold geometry for hypo-, normo- and hypertensive conditions, respectively. Since the pulse duplicator did not have an aortic root implemented but a straight connection downstream of the scaffold instead, the complex flow patterns during flow deceleration may not have developed as they would occur in a physiological aorta.

The evaluation of the high-speed camera acquisitions supports the comparability of scaffold geometries. The percentage value of the GOA did not change for a fixed pressure condition. For the hypertensive case with increased stroke volume, the GOA increased by 10% (cf. overlay in Figure 7) compared to other pressure conditions. The open-source, automated video analysis allowed for a reproducible evaluation of the GOA.

To further investigate the opening and closing over time, we used phase plots of GOA and flow. During the opening phase, when the flow is defined by the pump, the plots may enable an estimate of the scaffold’s parameters stiffness and elasticity. The scaffolds exhibited a similar opening curve in the phase plot from which we hypothesize that they had comparable stiffnesses. Since the same precursor material was used (theoretically yielding the same elasticity), we infer that the thickness was correctly quantified by the laser measurement during electrospinning. This was confirmed by SEM thickness measurements (not shown). The direct optical GOA measurement from high-speed video recordings provides cycle-resolved spatial information on opening dynamics that complements the ISO 5840 effective orifice area (EOA). Unlike Doppler echocardiography, it does not require acoustic access or assumptions about flow profiles, which makes it well suited to the optical inspection window available in standard pulse duplicator setups. GOA and EOA are correlated Adda et al. (2021).

The relationship between flow and GOA allows a further interpretation. To reach the hypertensive target pressures, stroke volume was increased by one third (from 60 mL to 80 mL), whereas maximum GOA increased by only about 10 %. By continuity, the flow rate through the valve is 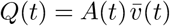,with orifice area *A* and mean transvalvular velocity 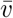,so that the stroke volume is 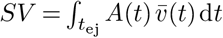 over the ejection period *t*_ej_. Since heart rate and systolic duration (and thus *t*_ej_) were identical in all conditions, the larger stroke volume had to be carried by a larger orifice, a higher velocity, or both. With GOA increasing by about 10 %, the mean transvalvular velocity must have increased by roughly 20 % (1.33*/*1.10 *≈* 1.2); most of the additional flow was therefore carried by a higher velocity rather than by a wider opening. Conversely, the hypo- and normotensive conditions, which used the same stroke volume, produced identical GOA waveforms despite 20 mmHg higher systolic and diastolic pressures in the normotensive case, indicating that leaflet opening was governed by flow rather than by the pressure level. Adjusting stroke volume and thus cardiac output to reach target pressures is common practice in in-vitro valve testing (e.g. Jahren et al., 2017). As a consequence, flow fields and leaflet stresses likely differ between pressure conditions even where GOA changes only slightly, a distinction that optical GOA tracking cannot resolve; flow-field imaging such as particle image velocimetry or flow magnetic resonance imaging would be required to characterize it directly. The small GOA differences between pressure conditions therefore most likely reflect the adjustment of stroke volume rather than a direct effect of pressure on leaflet mechanics. Overall, leaflet geometry had no systematic influence on the measured flow and pressure waveforms or on GOA under the experimental conditions studied.

### Limitations

The evaluation was restricted to four mandrel geometries representing a targeted subset of the parameter spaces of two analytical models; a comprehensive parametric sweep lies outside the scope of the current work. Bueno et al. (2026) systematically isolated individual geometric parameters in silicone valves and quantified leaflet strain fields by digital image correlation. Instead, the study at hand varies geometry at the level of the complete mandrel model and does not provide full-field strain data on the electrospun leaflets. Strain measurement on fibrous, porous electrospun leaflets under physiological flow remains technically challenging and is an important direction for future work to complement the hemodynamic GOA data reported here. The evaluation was limited to acute in-vitro hydrodynamic testing and does not address long-term scaffold durability, mechanical fatigue or cell-mediated remodeling, the endpoints that ultimately determine TEHV performance in-vivo. The mechanical properties of PCL materials have been extensively studied elsewhere, however. Leaflet coaptation was quantified by video recordings only and the test fluid was water rather than a blood analog; both factors limit the direct translatability of absolute regurgitation values. To further examine the influence of the microstructure on functionality testing and narrow the gap between characteristics in-vitro and in-vivo, the scaffold’s porosity may be reduced by a fibrin coating before testing. This and other coatings are currently being investigated by the authors.

A further limitation is that the pulse duplicator signals (pressure and flow) and the camera recordings used for GOA extraction were not synchronized by a common hardware trigger or timestamp. Instead, the two data streams were aligned by matching the start of forward flow (FFstart) to the rising edge of the GOA, since flow rate and GOA have been shown to share similar waveform shapes and maxima in-vitro (Zhang et al., 2018). Because this alignment was applied identically to all recordings, phase plots can be compared between scaffolds and pressure conditions, but the absolute phase lag between flow and GOA at valve opening cannot be determined and fine time-resolved comparisons between specific flow or pressure features and the GOA response should be avoided. After alignment, the residual timing uncertainty is on the order of the camera frame interval (6.7 ms at 150 fps). The distance between the upstream flow sensor and the valve does not add appreciably to this uncertainty: changes in flow rate propagate through the fluid-filled connection as pressure waves rather than at the fluid velocity, so that over the short, stiff section between sensor and valve the delay is well below one frame interval. At the same time, aligning the data on waveform features rather than on a hardware trigger makes the analysis independent of the sensor position and of setup-specific synchronization hardware, which facilitates transfer of the workflow to other pulse duplicator setups.

## Conclusion

Whole-valve electrospinning onto parametrized, 3D-printed mandrels enables the reproducible fabrication of electrospun PCL TEHV scaffolds that show physiological opening and closing kinematics across the full range of clinically relevant aortic pressure conditions. In contrast to other workflows, it highlights functionality testing of the electrospun scaffold material itself, preserving the fibrous microstructure and mechanical isotropy directly relevant to in-vivo remodeling. Mandrel geometry did not determine scaffold morphology nor the geometric orifice area. Flow and pressure waveforms remain broadly preserved across geometries for fixed boundary conditions, yet with a regurgitant fraction above ISO 5840 limits. These findings provide a reproducible evaluation framework for electrospun whole-valve scaffolds, supporting their continued development toward regenerative heart valve replacement.

## ACKNOWLEDGEMENTS

A.S., M.M. and C.V.C.B. acknowledge support from the Gravitation Program “Materials-Driven Regeneration” (MDR), funded by the Netherlands Organization for Scientific Research (NWO; grant no. 024.003.013). S.G., I.V.D.BV. and C.V.C.B. acknowledge funding from the Holland Hybrid Heart consortium, supported by the Dutch Research Agenda (NWA) of the Dutch Research Council (NWO) grant no. NWA.1518.22.049. H.D. acknowledges funding from the European Union’s Horizon Europe research and innovation programme under the Marie Skłodowska-Curie grant agreement No 101152298 (I3).

The authors would like to thank Sandra Loerakker and Justina Ghebryal for providing the initial implementation of the Hamid geometry and Marcel Rutten and Sandra Loerakker for the inspiring discussions.

## AUTHOR CONTRIBUTIONS

H.D.: Conceptualization, Methodology, Software, Investigation, Formal analysis, Data curation, Visualization, Funding acquisition, Project administration, Writing - original draft, Writing – review & editing. A.S.: Resources (3D printing, mandrel and scaffold preparation), Writing – review & editing. M.M.: Investigation (electrospinning, SEM imaging), Writing – review & editing. S.G.: Investigation (electrospinning, SEM imaging), Writing – review & editing. I.V.D.BV.: Investigation (electrospinning, SEM imaging), Writing – review & editing. C.V.C.B.: Supervision, Resources, Writing – review & editing.

All authors read and approved the final manuscript.

## DATA AVAILABILITY

Open-source data and code will be available at https://gitlab1.ptb.de/dillin01/functionaltehv.

## Supplementary Information

**Figure S1.**
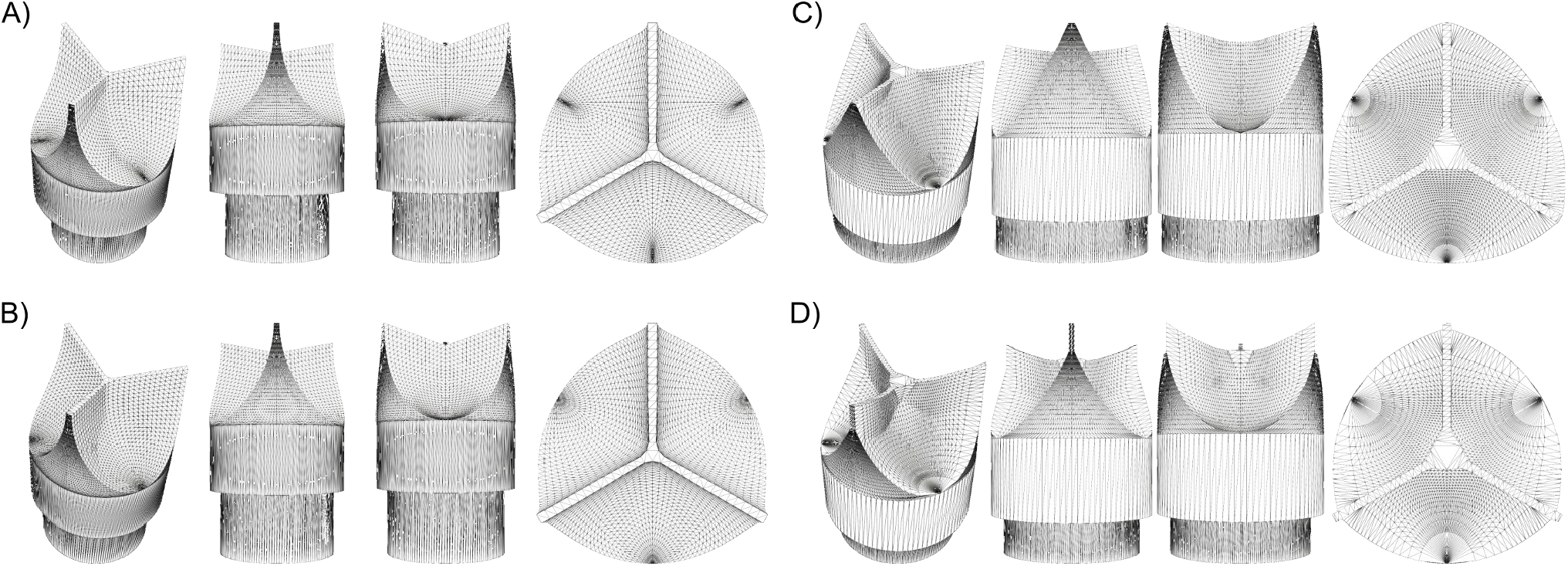
3D mesh models of the geometries used in the manuscript. A) Gulbulak *D* = 0, B) Gulbulak *D* = 0.5, C) Hamid *A* = 2.8, *B* = 1.4, C) Hamid *A* = 4.0, *B* = 2.4. Stereolithography files for 3D printing can be found in the Supplementary Material uploads.

**Figure S2.**
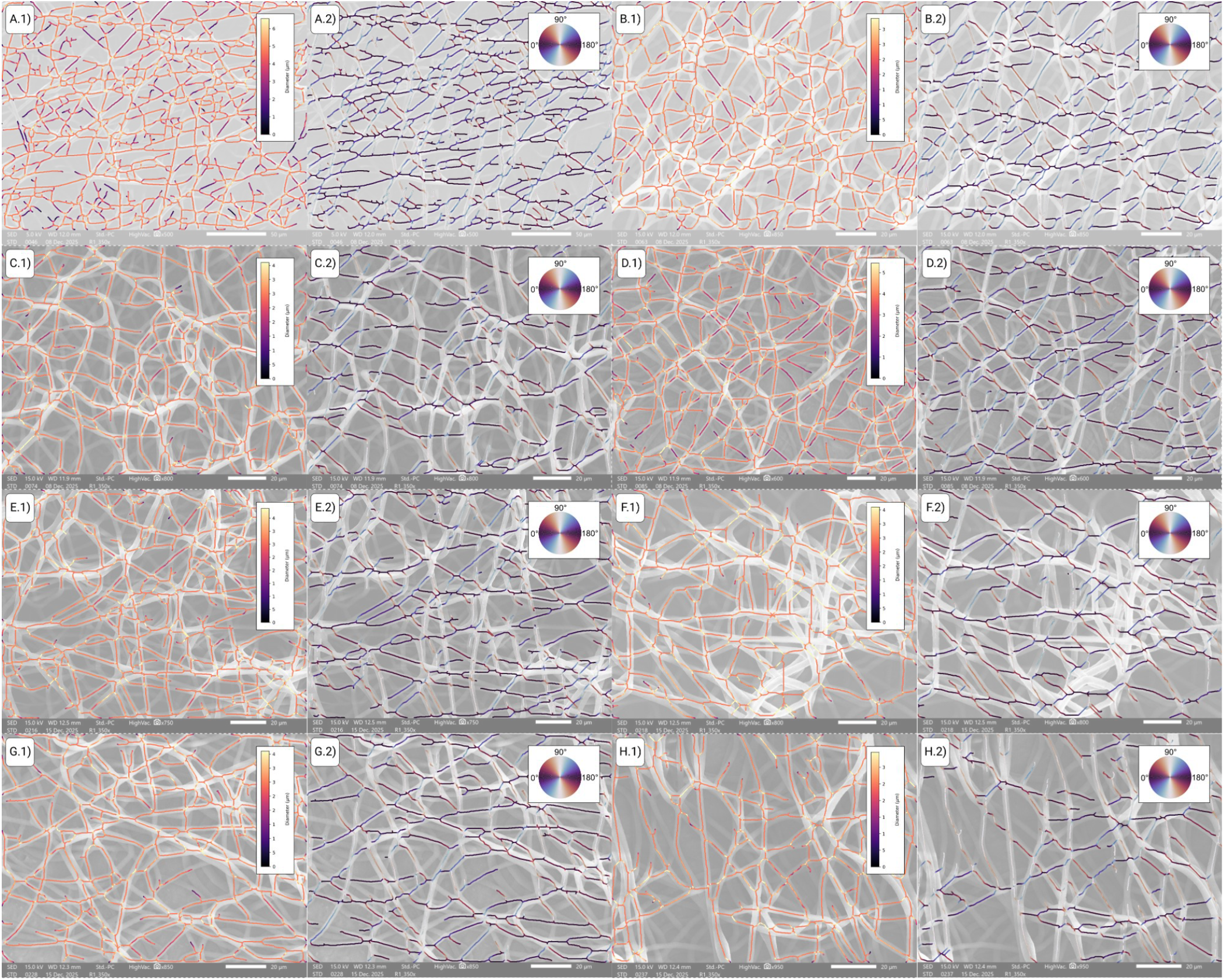
Visualization of automatic fiber parameter evaluation for scaffolds. Fiber diameters (.1) are homogeneous throughout the samples and approximately 2.5 µm. Fiber orientations (.2) show no preferred direction. N.B. the colormap shows vertical fibers in light colors which may limit their visibility. Scaffold numbers are A) #1.2, B) #1.5, C) #2.1, D) #2.4, E) #3.5, F) #3.6, G) #4.2, H) #4.3

**Figure S3.**
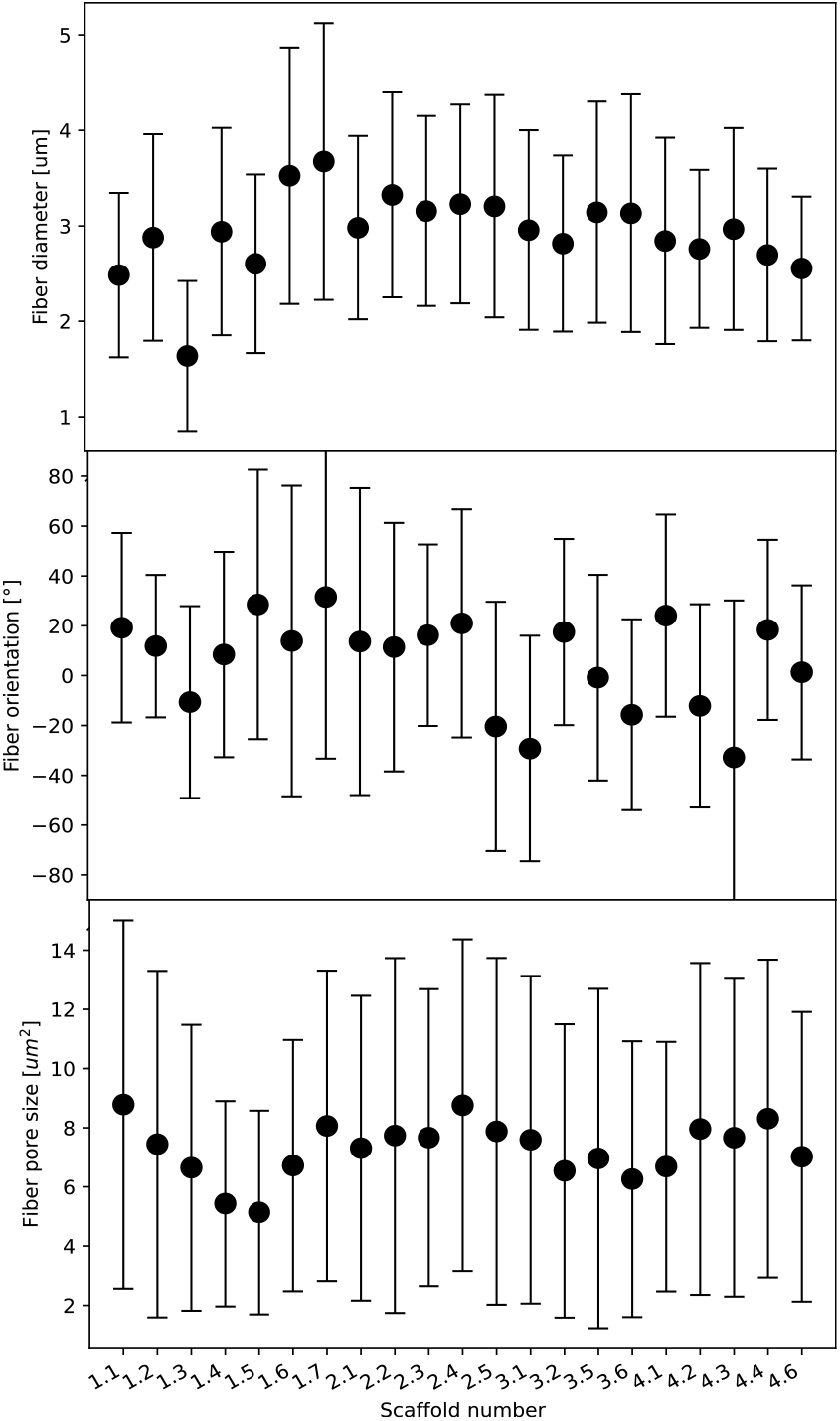
Evaluation of fiber parameters of the 21 electrospun scaffolds, sorted according to their batch. No trend is detected, indicating a stable process across batches. Top: Mean fiber diameter per scaffold is between 2.5 µm and 4 µm. Middle: Mean fiber orientation per scaffold is 0°, i.e. random orientation and no preferential alignment. Bottom: Mean fiber pore size per scaffold is 8 µm^2^. All evaluations have been performed automatically and may depend on the chosen segmentation settings.

**Figure S4.**
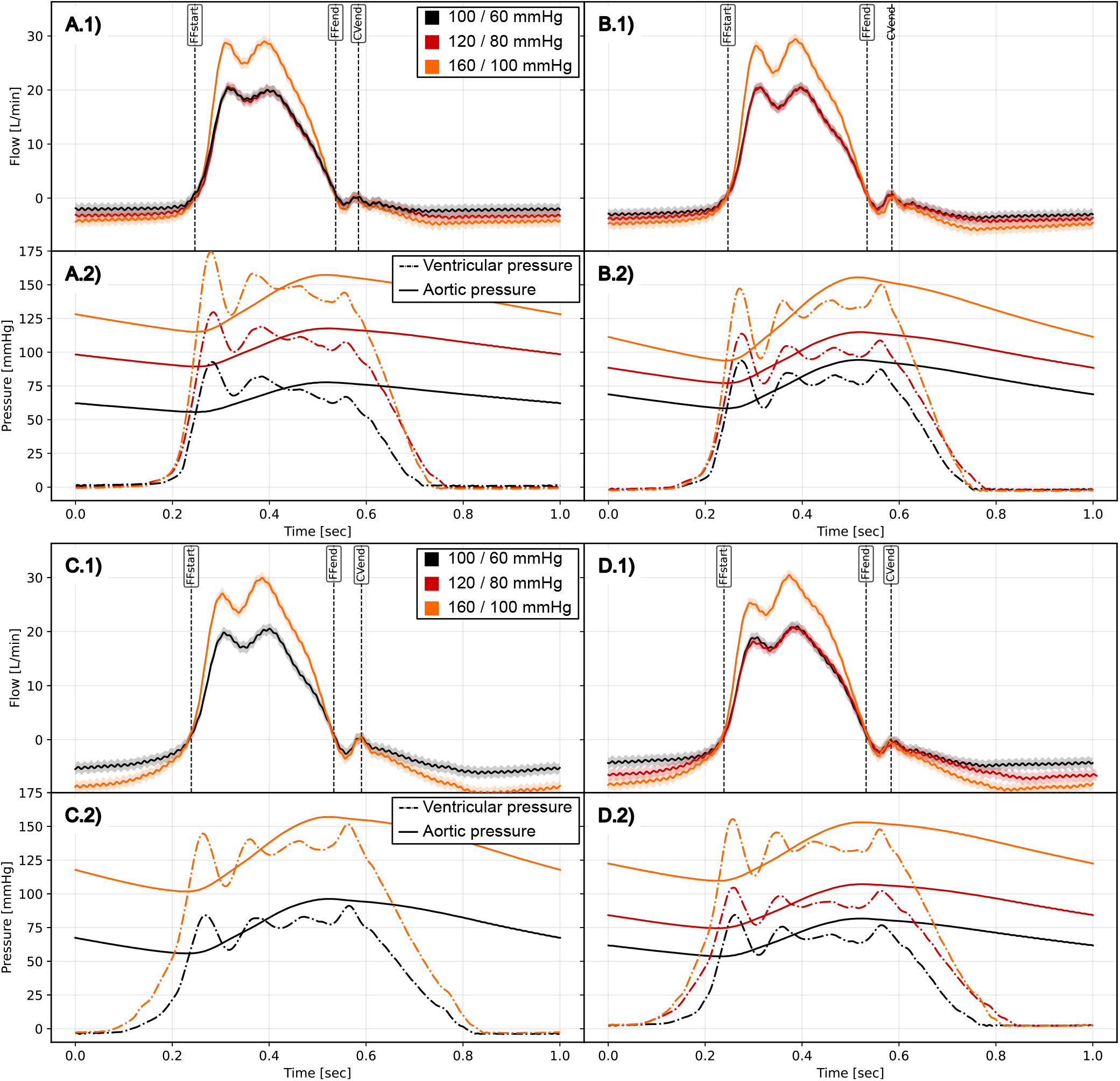
Geometries A) Gulbulak *D* = 0, B) Gulbulak *D* = 0.5, C) Hamid *A* = 2.8, *B* = 1.4, C) Hamid *A* = 4.0, *B* = 2.4 yielding flow (.1) and pressure curves (.2).

**Figure S5.**
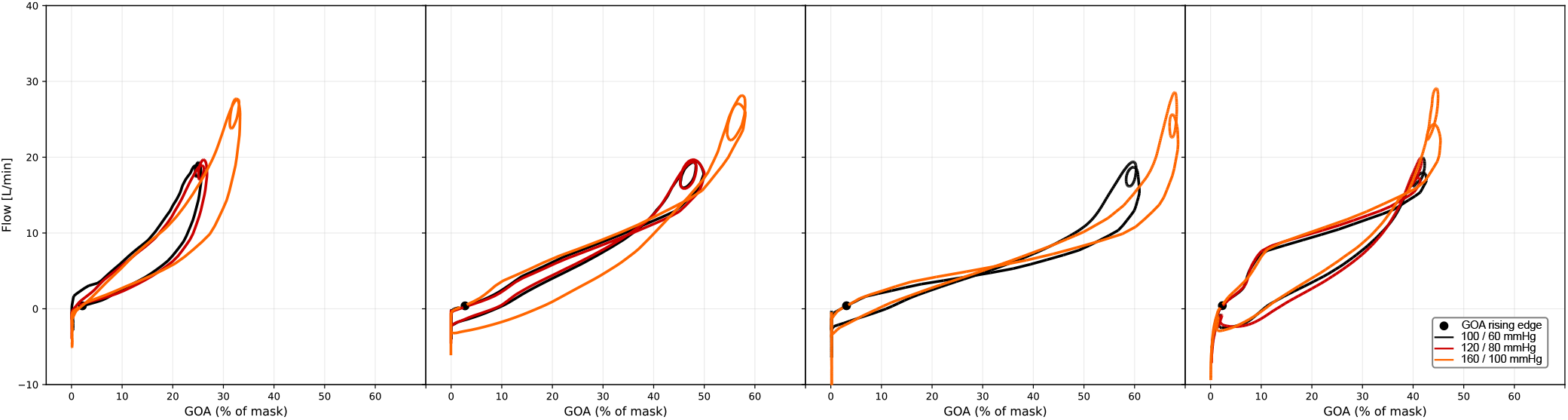
Phase plots GOA vs. flow for the different geometries. A) Gulbulak *D* = 0, B) Gulbulak *D* = 0.5, C) Hamid *A* = 2.8, *B* = 1.4, D) Hamid *A* = 4.0, *B* = 2.4. #3.6, #1.2, #2.1, #1.5

